# Determining the optimal SARS-CoV-2 mRNA vaccine dosing interval for maximum immunogenicity

**DOI:** 10.1101/2022.03.01.482592

**Authors:** Michael Asamoah-Boaheng, David M. Goldfarb, Martin A. Prusinkiewicz, Liam Golding, Mohammad Ehsanul Karim, Vilte Barakauskas, Nechelle Wall, Agatha N. Jassem, Ana Citlali Marquez, Chris MacDonald, Sheila F. O’Brien, Pascal Lavoie, Brian Grunau

**Affiliations:** Department of Emergency Medicine, University of British Columbia, Canada; Faculty of Medicine, Clinical Epidemiology, Memorial University of Newfoundland, Canada; Department of Pathology and Laboratory Medicine, University of British Columbia, Canada; Department of Pediatrics, University of British Columbia, Canada; Department of Obstetrics and Gynecology, University of British Columbia, Canada; Centre for Health Evaluation & Outcome Sciences, University of British Columbia, Canada; School of Population and Public Health, University of British Columbia, Canada; British Columbia Emergency Health Services, British Columbia, Canada; Public Health Laboratory, British Columbia Centre for Disease Control, British Columbia, Canada; Dalla Lana School of Public Health, University of Toronto, Toronto, Canada; Ontario Occupational Cancer Research Centre, Ontario, Canada; Canadian Blood Services, Canada; School of Epidemiology & Public Health, Universtiy of Ottawa, Ottawa, Canada

**Keywords:** SARS-CoV-2, vaccine dosing interval, spike total antibody concentrations, immunogenicity

## Abstract

**Objective:** Emerging evidence indicates that longer SARS-CoV-2 vaccine dosing intervals results in an enhanced immune response. However, the optimal vaccine dosing interval for achieving maximum immunogenicity is unclear.

**Methods:** This study included samples from adult paramedics in Canada who received two doses of either BNT162b2 or mRNA-1273 vaccines and provided blood samples 6 months (170 to 190 days) after the first vaccine dose. The main exposure variable was vaccine dosing interval (days), categorized as “*short*” (first quartile), “*moderate*” (second quartile), “*long*” (third quartile), and “*longest”* interval (fourth quartile). The primary outcome was total spike antibody concentrations, measured using the Elecsys SARS-CoV-2 total antibody assay. Secondary outcomes included: spike and RBD IgG antibody concentrations, and inhibition of angiotensin-converting enzyme 2 (ACE-2) binding to wild-type spike protein and several different Delta variant spike proteins. We fit a multiple log-linear regression model to investigate the association between vaccine dosing intervals and the antibody concentrations.

**Results:** A total of 564 adult paramedics (mean age 40 years, SD=10) were included. Compared to “short interval” (≤30 days), higher dosing interval quartiles (*moderate*: 31-38 days; *long*: 39-73 days and *longest*: ≥74 days) were all associated with increased Elescys spike total antibody concentration. Compared to the short interval, “*long*” and “*longest*” interval quartiles were associated with higher spike and RBD IgG antibody concentrations. Similarly, increasing dosing intervals increased inhibition of ACE-2 binding to viral spike protein, regardless of the vaccine type.

**Conclusion:** Increased mRNA vaccine dosing intervals longer than 30 days result in higher levels of circulating antibodies and viral neutralization when assessed at 6 months.

## Background

The coronavirus disease 2019 (COVID-19) caused by the SARS-CoV-2 virus was declared as a global pandemic on March 11, 2021 by the World Health Organization. SARS-CoV-2 vaccines were subsequently developed, with high efficacy for preventing short-term disease.^1,2^ The two mRNA vaccines, BNT162b2 (Pfizer) and mRNA-1273 (Moderna), were tested and have been approved for use as a two-dose schedule, administered 21 and 28 days apart, respectively.

Existing evidence indicates that extended vaccine dosing intervals for mRNA^3,4,5^ and Oxford-AstraZeneca^6^ vaccines enhance vaccine immunogenicity. However, while previous studies have investigated specific intervals, the exact optimal dosing interval for achieving maximum immunogenicity is unclear. Further, while extended dosing intervals may lead to an enhanced post-vaccine immune response, the trade-off is sub-total immunity in the between-dose period. Thus, the shortest interval to achieve the highest long-term immune response would be the optimal scenario.

For these reasons, we sought to investigate a range of dosing intervals and resultant immunogenicity (measured at 6-months post first vaccine), focusing on the wild-type and delta variant strains, as delta variants were the predominant strains at the time of blood sampling. We hypothesized that the benefits in increasing immunogenicity with increasing dosing intervals would plateau at a certain dosing interval, and thus sought to identify the shortest interval to achieve the maximum post-vaccine immune response.

## Methods

### Study design and data source

We used samples from the *COVID-19 Occupational Risks, Seroprevalence, and Immunity among Paramedics in Canada* (CORSIP) observational cohort study. CORSIP is a longitudinal prospective study examining the workplace risks and seroprevalence of SARS-CoV-2 exposure among paramedics (aged ≥ 19 years and older) working in the Canadian provinces of British Columbia, Ontario, Saskatchewan and Manitoba. Study participants provided responses to an administered questionnaire including past medical history (i.e., hypertension, diabetes, asthma, chronic lung disease, chronic heart disease, liver disease, malignancy, and immunosuppression), data pertaining to results and dates of vaccination and SARS-CoV-2 nucleic acid amplification tests (NAAT), and blood samples for serology testing. All samples used in this study were collected 170-190 days after the participant’s first vaccine dose. The CORSIP study was approved by the University of British Columbia and University of Toronto research ethics boards.

### Study participants

For this investigation we included samples from CORSIP participants who: (1) had two doses of either BNT162b2 or mRNA-1273 vaccines; and, (2) provided a blood sample 6-months ± 10 days (170-190 days) after the first vaccine. We excluded those with evidence of previous COVID-19 (defined as a positive NAAT COVID-19 test or presence of anti-nucleocapsid antibodies in the blood [Roche, IND, USA] ^**7**^) given the known differential impact on antibody responses post-vaccination.^8^ We also excluded cases for whom the second vaccine was >130 days after the first, in order to separate the vaccine and the blood collection date by >40 days, given the expected post-vaccine antibody surge.^6, 9^

### Serological Testing

All samples were tested with: (1) the Roche Nucleocapsid Elecsys Anti-SARS-Cov-2 (Roche, IND, USA) assay (to confirm eligibility); (2) the quantitative Roche Spike Elecsys Anti-SARS-Cov-2 S assay (Roche, IND, USA), which measures spike total antibody concentrations; (3) the V-PLEX COVID-19 Coronavirus Panel 2 IgG assay (Meso Scale discovery, MD, USA), which measures IgG to the SARS-CoV-2 spike and receptor binding domain (RBD) antigens; and (4) the V-PLEX SARS-CoV-2 Panel 19 ACE2 Kit (Meso Scale Discovery, MD, USA), which measures inhibition of angiotensin-converting enzyme 2 (ACE-2) binding to multiple delta (B.1.617.2) variant spike proteins.

### Outcome Measures

The primary outcome was spike total antibody concentrations, measured with the Elecsys assay. Secondary outcomes included: spike and RBD IgG antibody concentrations (measured with the V-PLEX assay), and inhibition of angiotensin-converting enzyme 2 (ACE-2) binding to wild-type spike protein and several different Delta (B.1.617.2) variant spike proteins (AY.1, AY.2, B.1.617.2/AY.3/AY.5/AY.6/AY.7/AY.14, B.1.617.2/AY.4, and AY.12).

### Statistical analysis

We performed statistical analyses with SPSS (IBM, USA) and R (V. 3.6.1) software. For participant characteristics we described categorical variables as counts (with percentages) and continuous variables as mean (with standard deviation [SD]) or median (with interquartile range [IQR]) values. To visualize the trend of the antibody response over the range of dosing intervals, we created: (1) error plots of the means (with 95% confidence intervals) of outcomes values; and, (2) created scatter-plots with cubic spline curves (with 95% confidence intervals).^10^

To investigate the relationship between vaccine dosing intervals and antibody outcomes, we divided participants into vaccine dosing interval quartiles: “*short interval*” (first quartile), “*moderate interval*” (second quartile), “*long interval*” (third quartile), and “*longest interval*” (fourth quartile). We compared characteristics and outcomes between quartiles using ANOVA to test for differences in the mean ages of participants, Chi-square test to test for differences between categorical variables, and the Kruskal Wallis H test for the difference between outcome measures.

We fit a multiple log-linear regression model to estimate the association between the dosing interval quartiles (with the first quartile as the reference) and both the primary and secondary outcome variables, adjusted for type of vaccine, sex, age, and medical history (each past medical history category was modeled as a binary variable). As sensitivity analyses, we repeated the primary analyses incorporating an interaction term with dosing interval category and vaccine type. We also repeated the primary analysis within subgroups based on vaccine type.

## Results

The study included a total of 564 adult paramedics, with a mean age of 40 years; 253 (45%) were female sex at birth and 79 (14%) were from Asia and other ethnic groups (Table 1). Overall, 469 (83%) received two doses of the BNT162b2 vaccine and 95 (17%) received two doses of the mRNA-1273 SARS-CoV-2 vaccine.

**Table 1:**
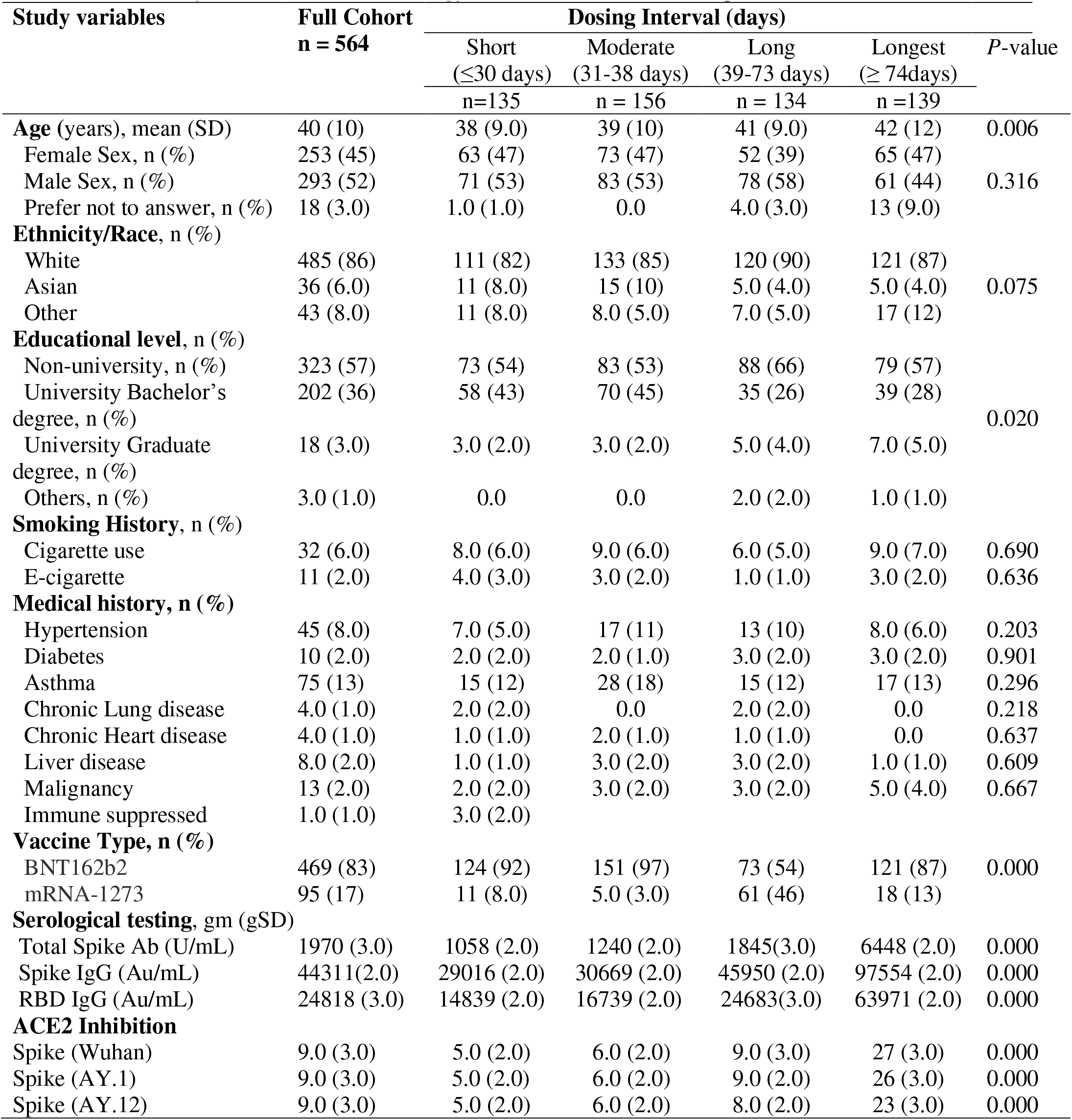
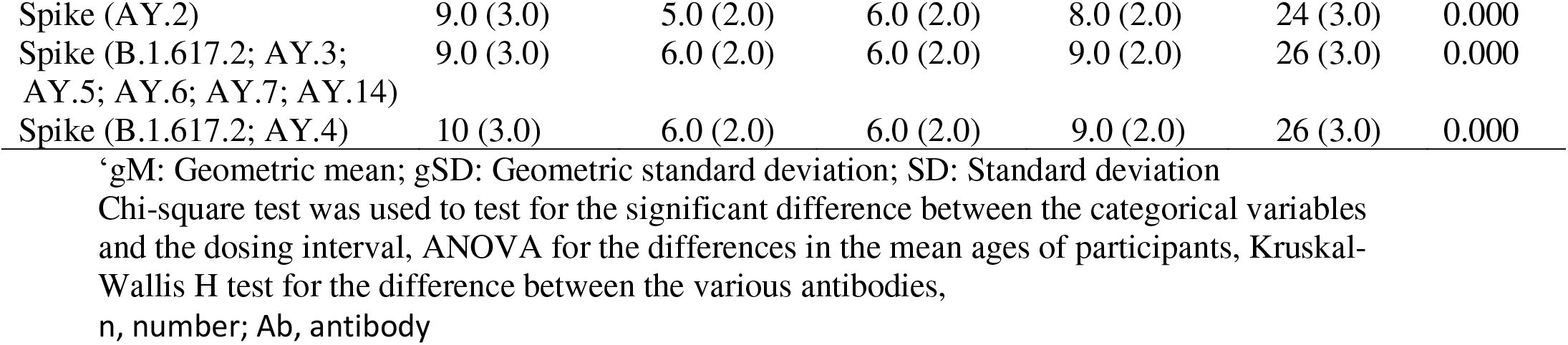
Study characteristics and serology results of the 6 months sample

The median vaccine dosing interval was 38 days (IQR 31-73), and thus cases were classified “short interval” (≤30 days), “moderate interval” (31-38 days), “long interval” (39-73 days), and “longest interval” (≥74 days). Table 1 shows participant characteristics, which were similar across dosing interval quartiles. All outcome measures increased with each vaccine dosing interval quartile. Figures 1, 2, 3 and Supplementary figures S1, S3, S5, S7, S9, S10, S12-S13 show mean error plots and spline graphs of outcome measures over a range of vaccine dosing intervals, demonstrating an increase in all measures of immunogenicity with increasing dosing intervals. A similar relationship was seen among individuals who were fully vaccinated with either BNT162b2 and mRNA-1273 vaccines (see Supplementary Figures S2, S4, S6, S8, S11).

**Figure 1:**
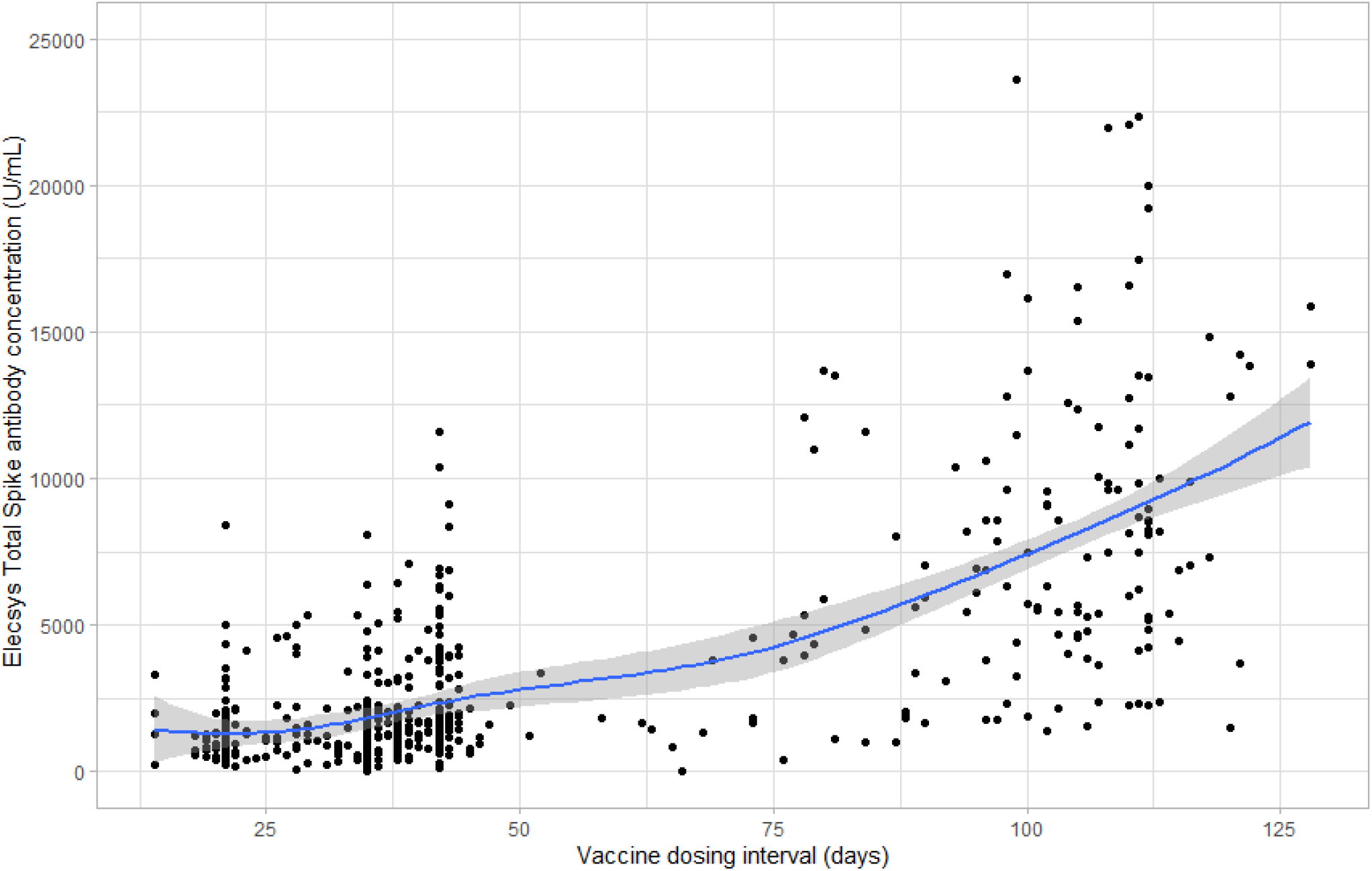
Scatter plot of the Elecsys spike total antibody concentration (U/mL) against the vaccine dosing interval (days)

**Figure 2:**
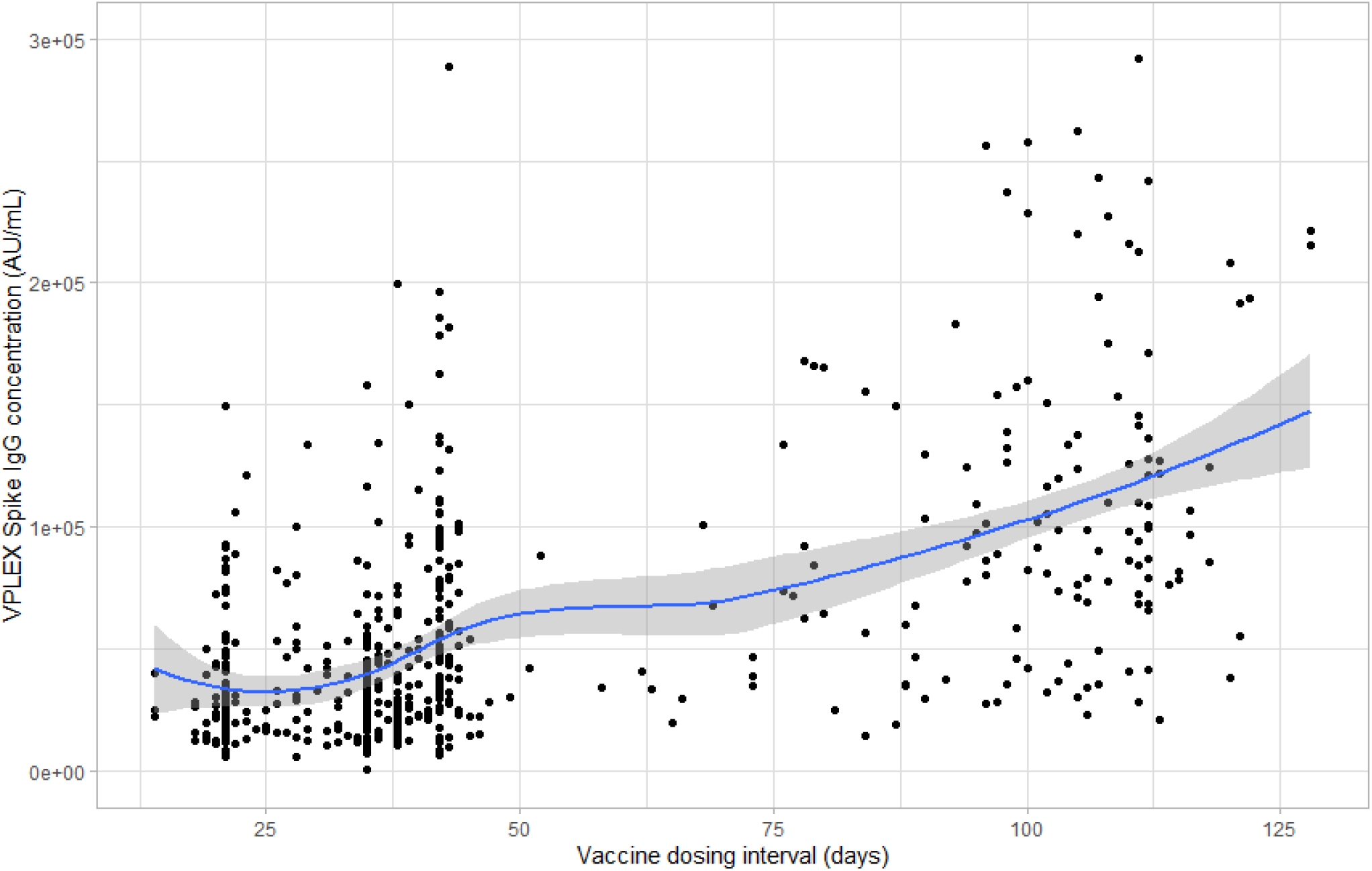
Scatter plot of the V-PLEX spike IgG antibody concentration (AU/mL) against the vaccine dosing interval (days)

**Figure 3:**
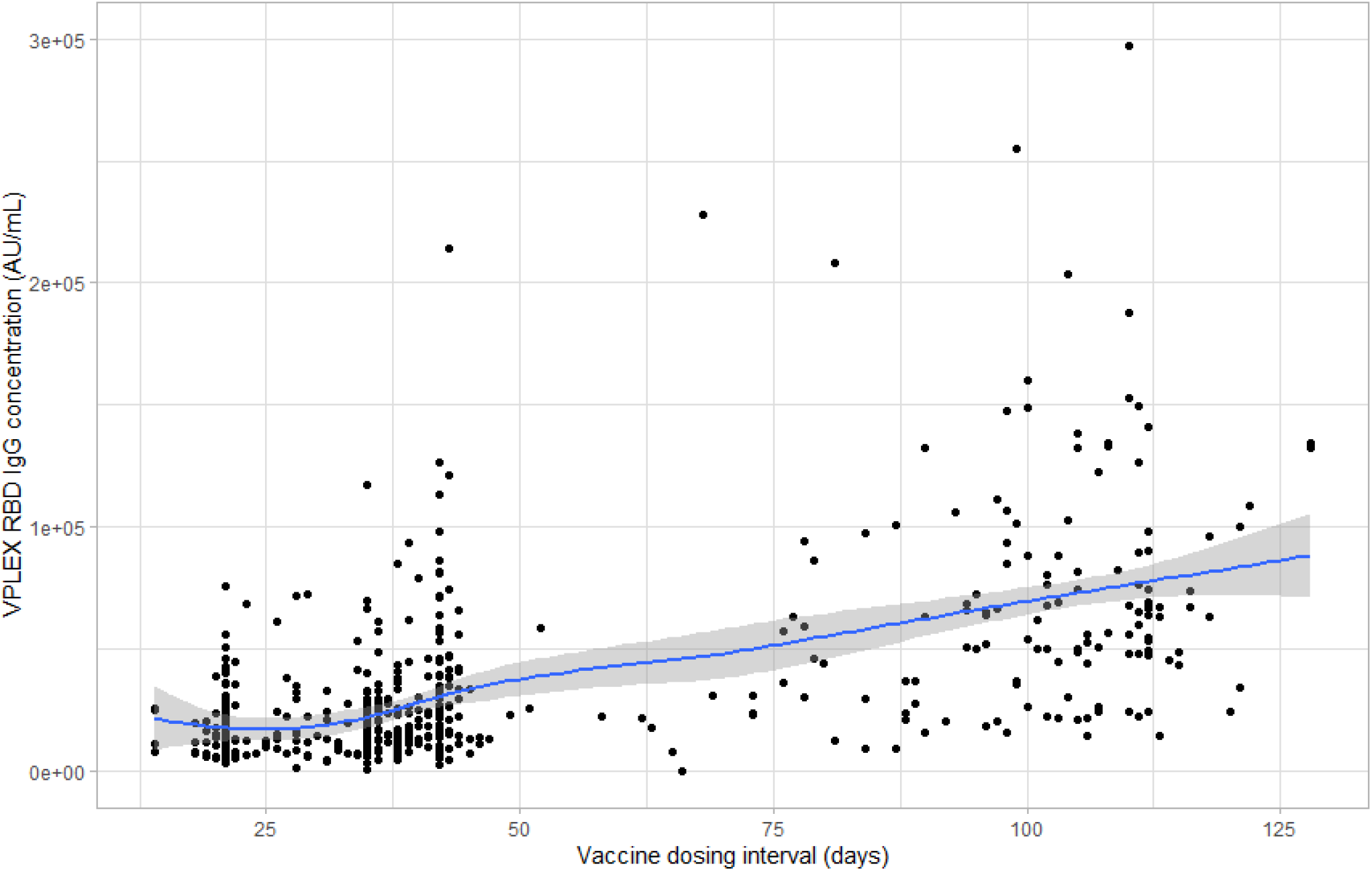
Scatter plot of the V-PLEX RBD antibody concentration (AU/mL) against the vaccine dosing interval (days)

The primary regression analyses demonstrated that, with reference to the first quartile, higher quartiles were all associated with increased log Elecsys total spike antibody concentrations (Table 2). Models examining secondary outcomes of V-PLEX spike and RBD IgG showed that “long” and “longest” quartiles, with reference to the first quartile, were associated with significantly increased log antibody concentrations. Models examining secondary outcomes of ACE-2 inhibition to the wild type spike protein and Delta strain spike variant proteins showed similar results (see Table 3).

**Table 2:** Multivariate analysis (Multiple log-linear regression) of the association between vaccine dosing intervals and antibody

| Dosing intervals | Model 1 |  | Model 2 |  | Model 3 |  |
| --- | --- | --- | --- | --- | --- | --- |
| | Coeff ($\beta$) | 95% CI | Coeff ($\beta$) | 95% CI | Coeff ($\beta$) | 95% CI |
| *Moderate* (31-38 days) | 0.19 | (0.030, 0.34) ** | 0.09 | (-0.070, 0.25) | 0.17 | (-0.003, 0.34) |
| *Long* (39-73 days) | 0.33 | (0.16, 0.50) ** | 0.31 | (0.13, 0.48) ** | 0.35 | (0.15, 0.54) ** |
| *Longest* ($\geq 74$days) | 0.90 | (0.74, 1.10) ** | 1.27 | (1.11, 1.43) ** | 1.52 | (1.34, 1.69) ** |

*Model 1:* Outcome variable in model 1 is Elecsys Spike total Antibody Concentration; *Model 2:* Outcome variable in model 2 is V-PLEX Spike IgG Antibody Concentration; *Model 3:* Outcome variable in model 3 is V-PLEX RBD Antibody Concentration.
\*\*indicates model is significant at $p < 0.05$
All models were adjusted for type of vaccine administered, age, “sex at birth”, and comorbidities (hypertension, diabetes, asthma, chronic lung diseases, heart diseases, kidney diseases, liver diseases, cancer, and immune suppressed).
Low dosing interval ( $< 31$ days) was set as a reference category

**Table 3:**
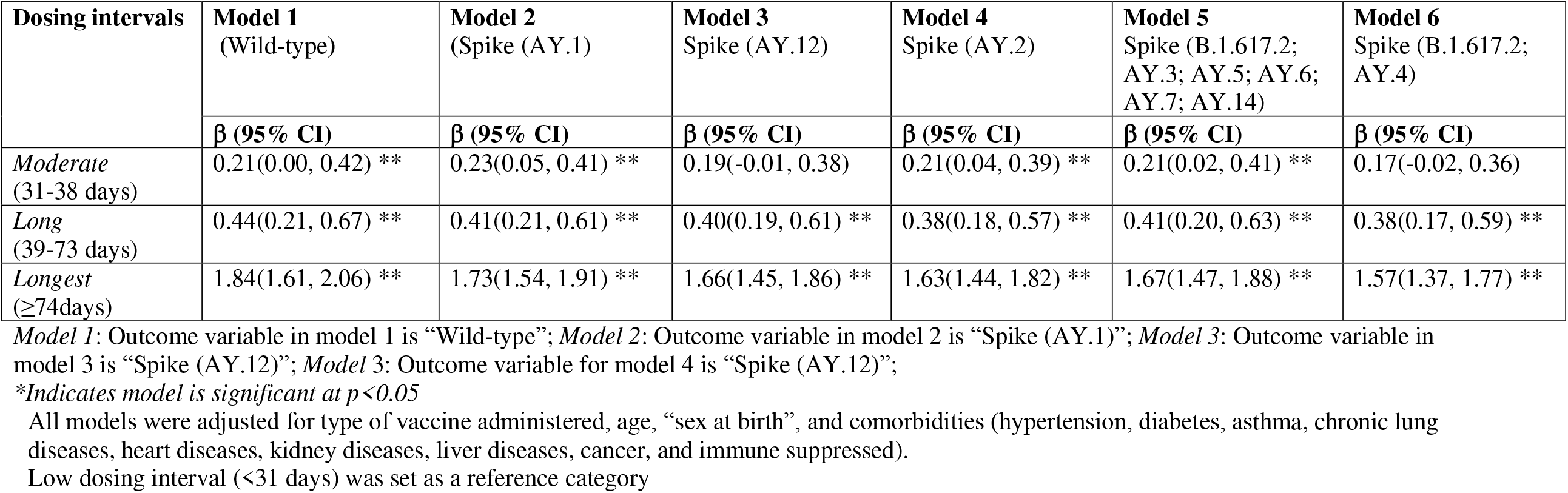
Multivariate analysis (Multiple log-linear regression) of the association between vaccine dosing intervals and secondary antibody outcomes

| Dosing intervals | Model 1<br>(Wild-type) | Model 2<br>(Spike (AY.1)) | Model 3<br>Spike (AY.12) | Model 4<br>Spike (AY.2) | Model 5<br>Spike (B.1.617.2;<br>AY.3; AY.5; AY.6;<br>AY.7; AY.14) | Model 6<br>Spike (B.1.617.2;<br>AY.4) |
| --- | --- | --- | --- | --- | --- | --- |
| | $\beta$ (95% CI) | $\beta$ (95% CI) | $\beta$ (95% CI) | $\beta$ (95% CI) | $\beta$ (95% CI) | $\beta$ (95% CI) |
| *Moderate*<br>(31-38 days) | 0.21(0.00, 0.42) ** | 0.23(0.05, 0.41) ** | 0.19(-0.01, 0.38) | 0.21(0.04, 0.39) ** | 0.21(0.02, 0.41) ** | 0.17(-0.02, 0.36) |
| *Long*<br>(39-73 days) | 0.44(0.21, 0.67) ** | 0.41(0.21, 0.61) ** | 0.40(0.19, 0.61) ** | 0.38(0.18, 0.57) ** | 0.41(0.20, 0.63) ** | 0.38(0.17, 0.59) ** |
| *Longest*<br>($\geq 74$days) | 1.84(1.61, 2.06) ** | 1.73(1.54, 1.91) ** | 1.66(1.45, 1.86) ** | 1.63(1.44, 1.82) ** | 1.67(1.47, 1.88) ** | 1.57(1.37, 1.77) ** |

\*Indicates model is significant at $p < 0.05$
All models were adjusted for type of vaccine administered, age, “sex at birth”, and comorbidities (hypertension, diabetes, asthma, chronic lung diseases, heart diseases, kidney diseases, liver diseases, cancer, and immune suppressed).
Low dosing interval (<31 days) was set as a reference category

**Table 4:**
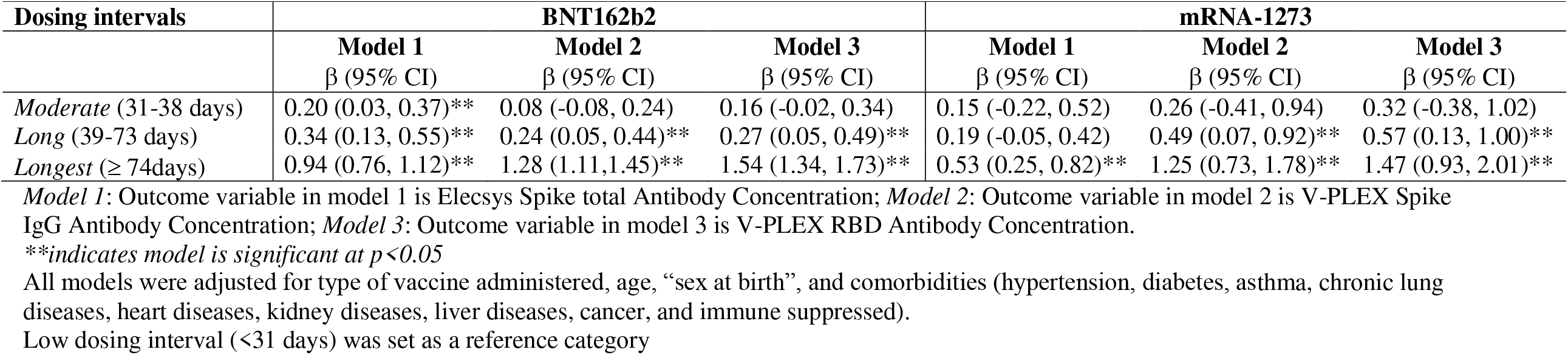
Multiple log-linear regression between dosing intervals and antibody response by vaccine type

In the sensitivity analysis, we introduced an interaction term of vaccine type and the vaccine dosing interval into all regression models; however, none were significant (Tables S1 and S2). We repeated the models in subgroups defined by vaccine type; although limited by smaller sample sizes, results were similar.

## Discussion

We investigated 6-month immunogenicity among 564 adults who received SARS-COV-2 mRNA vaccines, to identify the vaccine dosing interval for achieving maximum immunogenicity. We found that longer vaccine dosing intervals resulted in higher measures of immunogenicity 6-months post-vaccination, including antibodies against spike protein and its RBD domain, as well as surrogate measures of viral neutralization. Whereas we hypothesized that the rise in immune measures would plateau with a certain vaccine dosing interval, antibody levels continued to rise with extended vaccine intervals up to 130 days. This data will assist clinicians and policy-makers by demonstrating that increasing vaccine dosing intervals (up to 130 days) results in corresponding increased 6-month immunogenicity.

Our data demonstrated a continuous increasing trend of immunogenicity as the vaccine dosing interval increased, which was consistent with all immune measures. Our regression models were consistent. In deciding the optimal vaccine dosing interval, there is a trade-off between decreased immunity between the first and second dose with longer dosing intervals, and enhanced immunity in the post-second dose period. We had hoped to identify a plateau in the immunogenicity measures with increasing dosing intervals—however this was not apparent.

Instead, public health providers need to balance these two competing priorities, the balance of which may change based on the incidence of COVID-19 in the community, and the risk of severe disease in individual patients. Of additional consideration is the availability of booster (third) vaccine dosing—i.e. if readily available then decreased immunity in the post-second dose period among those with short vaccine dosing intervals can be addressed by a booster dose.^11, 12^

Community level-immunity may also play a role in vaccine dosing interval decisions. Previous modeling studies have demonstrated that extending SARS-CoV-2 vaccine dosing intervals results in reduced cumulative mortality. This is due to earlier access to the first vaccine dose, even without considering the long-term benefit of post-second dose enhanced immunogenicity.^13,14^ Global-level immunity and vaccine distribution also deserve consideration. While booster doses may serve as a method to augment waning immunity in the months after the second mRNA dose, this diverts vaccine supplies to low and middle incomes where only 10.4% of the population has received at least 1 dose of the vaccine.^15^ Lengthening between-dose intervals may result in more robust longer-term immunity, thus allowing for delayed booster doses, and re-direction of vaccine supplies to under-served regions.

Our results are consistent with recently released (pre-peer review) data, showing that a 7-8 week interval between doses improved vaccine effectiveness in comparison to a 3-4 week interval.^16^ Our data also corroborates immunogenicity studies that compared an extended dosing interval schedule of the BNT162b2 vaccine to the standard schedule (3 weeks).^5,17^ A recent study by Parry et al (2021)^18^ found that delaying or extending the timing of the second dose of the BNT162b2 vaccine up to 12 weeks generated higher antibody levels with 3.5-fold higher peak antibody values than the standard 3-week interval. Additionally, higher two dose vaccine effectiveness was observed when more than 6 weeks dosing interval was adopted between BNT162b2 doses compared to the standard schedule.^19^ Similar findings were found in studies conducted by Moghadas et al.^15^ and Payne et al.^20^

Some jurisdictions such as the United States administer the second dose of the SAR-COV-2 vaccines by the stipulated 3-or 4-week schedules for Pfizer-BioNTech and Moderna vaccines, respectively. Few countries, including the United Kingdom (UK) and parts of Canada have approved guidelines for delaying the second dose schedules of the vaccines to up to 12 and 16 weeks respectively ^21, 22^. In contrast, the world health organization (WHO) has recently recommended an interval of at least 6 weeks between same mRNA vaccines.^23^ The existence of these varying dosing interval strategies raises some concerns as to which dosing interval is optimal in achieving maximal immunogenicity. Our study sought to investigate this phenomenon to identify the minimum optimal threshold to improve immunity against COVID-19; we found that increasing dosing intervals resulted in a continual increase in immunogenicity measures at 6 months.

## Limitations

This was an observational study, and it is possible that unaccounted for confounders affected our results. We measured antibody concentrations at 6-months, as we believed that immunogenicity at a standard time interval after the beginning of the vaccination series is the most clinically relevant end-point. Although we excluded samples for which the second vaccine-to-blood collection interval was <40 days, due to the expected post-vaccination antibody surge, it is possible that proximity of the second dose to our blood collection time affected the results. We excluded cases with previous COVID-19 using nucleocapsid serological testing—which, although a high-performance test ^7^, may have mis-classified samples. Our study participants consist of paramedics that were primarily mid-aged and white, and thus these data may not be generalizable to other groups. Although antibody levels have been correlated with vaccine effectiveness^24^, we did not examine clinical endpoints such as break-through infections or severe disease.

## Conclusion

Extending the interval between SARS-CoV-2 mRNA vaccine doses from 31 to 123 days is associated with a continuous increase in immunogenicity when measured at 6-months.

## Reference

1. Polack FP, Thomas SJ, Kitchin N, Absalon J, Gurtman A, Lockhart S, Perez JL, Pérez Marc G, Moreira ED, Zerbini C, Bailey R, Swanson KA, Roychoudhury S, Koury K, Li P, Kalina W V, Cooper D, Frenck RW, Hammitt LL, Türeci Ö, Nell H, Schaefer A, Ünal S, Tresnan DB, Mather S, Dormitzer PR, Şahin U, Jansen KU, Gruber WC, C4591001 Clinical Trial Group. Safety and Efficacy of the BNT162b2 mRNA Covid-19 Vaccine. N Engl J Med. 2020;383(27):2603–2615. doi:10.1056/NEJMoa2034577.

2. Baden LR, El Sahly HM, Essink B, Kotloff K, Frey S, Novak R, Diemert D, Spector SA, Rouphael N, Creech CB, McGettigan J, Khetan S, Segall N, Solis J, Brosz A, Fierro C, Schwartz H, Neuzil K, Corey L, Gilbert P, Janes H, Follmann D, Marovich M, Mascola J, Polakowski L, Ledgerwood J, Graham BS, Bennett H, Pajon R, Knightly C, Leav B, Deng W, Zhou H, Han S, Ivarsson M, Miller J, Zaks T, COVE Study Group. Efficacy and Safety of the mRNA-1273 SARS-CoV-2 Vaccine. N Engl J Med. 2021;384(5):403–416. doi:10.1056/NEJMoa2035389.

3. Payne RP, Longet S, Austin JA, Skelly DT, Dejnirattisai W, Adele S, Meardon N, Faustini S, Dunachie S, et al. Immunogenicity of standard and extended dosing intervals of BNT162b2 mRNA vaccine. Cell. October 2021. doi:10.1016/j.cell.2021.10.011.

4. Grunau B, Goldfarb DM, Asamoah-Boaheng M, Golding L, Kirkham TL, Demers PA, Lavoie PM. Immunogenicity of Extended mRNA SARS-CoV-2 Vaccine Dosing Intervals. JAMA. December 2021. doi:10.1001/jama.2021.21921.

5. Grunau B, Asamoah-Boaheng M, Lavoie PM, Karim ME, Kirkham TL, Demers PA, Barakauskas V, Marquez AC, Jassem AN, O’Brien SF, Drews SJ, Haig S, Cheskes S, Goldfarb DM. A Higher Antibody Response Is Generated With a 6-to 7-Week (vs Standard) Severe Acute Respiratory Syndrome Coronavirus 2 (SARS-CoV-2) Vaccine Dosing Interval. Clin Infect Dis. November 2021. doi:10.1093/cid/ciab938.

6. Voysey M, Costa Clemens SA, Madhi SA, Weckx LY, Folegatti PM, Aley PK, Angus B, Baillie VL, Zuidewind P, et al. Single-dose administration and the influence of the timing of the booster dose on immunogenicity and efficacy of ChAdOx1 nCoV-19 (AZD1222) vaccine: a pooled analysis of four randomised trials. Lancet. 2021;397(10277):881–891. doi:10.1016/S0140-6736(21)00432-3.

7. Ainsworth M, Andersson M, Auckland K, Baillie JK, Barnes E, Beer S, Beveridge A, Bibi S, Young RK, et al. Performance characteristics of five immunoassays for SARS-CoV-2: a head-to-head benchmark comparison. Lancet Infect Dis. 2020;20(12):1390–1400. doi:10.1016/S1473-3099(20)30634-4.

8. Ebinger JE, Fert-Bober J, Printsev I, Wu M, Sun N, Prostko JC, Frias EC, Stewart JL, Van Eyk JE, Braun JG, Cheng S, Sobhani K. Antibody responses to the BNT162b2 mRNA vaccine in individuals previously infected with SARS-CoV-2. Nat Med. 2021;27(6):981–984. doi:10.1038/s41591-021-01325-6.

9. Doria-Rose N, Suthar MS, Makowski M, O’Connell S, McDermott AB, Flach B, Ledgerwood JE, Mascola JR, Graham BS, Lin BC, O’Dell S, Schmidt SD, Widge AT, Edara V-V, Anderson EJ, Lai L, Floyd K, Rouphael NG, Zarnitsyna V, Roberts PC, Makhene M, Buchanan, W, Luke CJ, Beigel JH, Jackson LA, Neuzil KM, Bennett H, Leav B, Albert J, Kunwar P. Antibody Persistence through 6 Months after the Second Dose of mRNA-1273 Vaccine for Covid-19, New England Journal of Medicine. 2021, 384(23): 1–3

10. Gauthier J, Wu Q V., Gooley TA. Cubic splines to model relationships between continuous variables and outcomes: a guide for clinicians. Bone Marrow Transplant. 2020;55(4):675–680. doi:10.1038/s41409-019-0679-x.

11. An Advisory Committee Statement (ACS) National Advisory Committee on Immunization (NACI). Guidance on booster COVID-19 vaccine doses in Canada-Update December 3, 2021. 2021. https://www.canada.ca/content/dam/phac-aspc/documents/services/immunization/national-advisory-committee-on-immunization-naci/guidance-booster-covid-19-vaccine-doses/guidance-booster-covid-19-vaccine-doses.pdf. Accessed on February 7, 2022.

12. World Health Organization (WHO). Interim statement on booster doses for COVID-19 vaccination: update 4 October 2021. https://www.who.int/news/item/04-10-2021-interim-statement-on-booster-doses-for-covid-19-vaccination. Accessed on February 7, 2022.

13. Romero-Brufau S, Chopra A, Ryu AJ, Gel E, Raskar R, Kremers W, Anderson KS, Subramanian J, Krishnamurthy B, Singh A, Pasupathy K, Dong Y, O’Horo JC, Wilson WR, Mitchell O, Kingsley TC. Public health impact of delaying second dose of BNT162b2 or mRNA-1273 covid-19 vaccine: simulation agent based modeling study. BMJ. May 2021:n1087. doi:10.1136/bmj.n1087.

14. Moghadas SM, Vilches TN, Zhang K, Nourbakhsh S, Sah P, Fitzpatrick MC, Galvani AP. Evaluation of COVID-19 vaccination strategies with a delayed second dose. Read AF, ed. PLOS Biol. 2021;19(4):e3001211. doi:10.1371/journal.pbio.3001211.

15. Our World in Data. Coronavirus (COVID-19) Vaccinations. https://ourworldindata.org/covid-vaccinations. Accessed February 8, 2022.

16. Skowronski DM, Setayeshgar S, Febriani Y, Ouakki M, Zou M, Talbot D, et al. Two-dose SARS-CoV-2 vaccine effectiveness with mixed schedules and extended dosing intervals: test-negative design studies from British Columbia and Quebec, Canada, BMJ, 2021; 1–28. https://doi.org/10.1101/2021.10.26.21265397

17. Grunau B, Goldfarb DM, Asamoah-Boaheng M, Golding L, Kirkham TL, Demers PA, Lavoie PM. Immunogenicity of extended mRNA SARS-CoV-2 vaccine dosing intervals. JAMA. 2021 Dec 3, 2021. doi: 10.1001/jama.2021.21921

18. Parry H, Bruton R, Stephens C et al. Extended interval BNT162b2 vaccination enhances peak antibody generation in older people. medRxiv. 2021. doi:10.1101/2021.05.15.21257017.

19. Amirthalingam G, Bernal JL, Andrews NJ, Whitaker H, Gower C, Stowe J, Tessier E, Subbarao S, Ireland G, Baawuah F, Linley E, Warrener L, O’Brien M, Whillock C, Moss P, Ladhani SN, Brown KE, Ramsay ME. Serological responses and vaccine effectiveness for extended COVID-19 vaccine schedules in England. Nat Commun. 2021 Dec 10;12(1):7217. doi: 10.1038/s41467-021-27410-5. PMID: 34893611; PMCID: PMC8664823.

20. Payne RP, Longet S, Austin JA, Skelly DT et al. Immunogenicity of standard and extended dosing intervals of BNT162b2 mRNA vaccine, Cell. 2021; 184: 5699–5714

21. Covid-19 second-stage vaccinations to be delayed across UK. Guardian 2020 Dec 30. https://www.theguardian.com/world/2020/dec/30/covid-19-second-stage-nhs-vaccinations-delayed-across-uk. Accessed on January 19, 2022.

22. Department of Health and Social Care. Optimising the COVID-19 vaccination programme for maximum short-term impact. Updated January 26, 2021. https://www.gov.uk/government/publications/prioritising-the-first-covid-19-vaccine-dose-jcvi-statement/optimising-the-covid-19-vaccination-programme-for-maximum-short-term-impact. Accessed on January 19, 2021.

23. World Health Organization (WHO). An Advisory Committee Statement (ACS) National Advisory Committee on Immunization (NACI) Interim guidance on booster COVID-19 vaccine doses in Canada. Published October 29, 2021. https://www.canada.ca/content/dam/phacaspc/documents/services/immunization/national-advisory-committee-on-immunization-naci/recommendations-use-covid-19-vaccines/statement-guidance-booster-doses/statement-guidance-booster-doses.pdf. Accessed on January 19, 2021.

24. Earle KA, Ambrosino DM, Fiore-Gartland A, Goldblatt D, Gilbert PB, Siber GR, Dull P, Plotkin SA. Evidence for antibody as a protective correlate for COVID-19 vaccines. Vaccine. 2021;39(32):4423–4428. doi:10.1016/j.vaccine.2021.05.063.

